# C1Q-associated adaptive myeloid remodelling accompanies early response to BCMA CAR-T therapy in multiple myeloma

**DOI:** 10.64898/2026.08.23.746487

**Authors:** Shubing Wang, Qian Wang, Yan-Ruide Li, Song Li

**Author notes:** **Additional Information correspondence statement** Yan-Ruide Li, Ph.D., Department of Microbiology, Immunology & Molecular Genetics, University of California, Los Angeles, Los Angeles, CA 90095, USA., Song Li, Ph.D., Department of Bioengineering, University of California, Los Angeles, Los Angeles, CA 90095, USA.

## Abstract

BCMA-directed chimeric antigen receptor T cells induce deep responses in multiple myeloma, yet the immune ecology accompanying early response remains incompletely resolved. We reanalysed 171,971 single-cell transcriptomes from 25 peripheral-blood and bone-marrow specimens from ten patients. At day 30, responders showed concordant enrichment of C1Q and IFN-γ programmes in blood and marrow myeloid pseudobulk profiles. Antigen-presentation genes were enriched in blood, whereas TGF-β and hypoxia programmes were depleted in responding marrow. Cholesterol-efflux genes were not enriched in responders or after treatment. A composite C1Q-cholesterol score showed nominal associations with response and CD8 dysfunction in selected compartments, but none survived study-wide correction. The pathway results support an adaptive, antigen-presenting C1Q-associated programme rather than a uniformly suppressive C1Q macrophage model. This state-contingent interpretation of early myeloid remodelling requires prospective, patient-level validation before biomarker or causal claims are warranted.

## INTRODUCTION

BCMA-directed chimeric antigen receptor (CAR) T-cell therapy produces high response rates in relapsed or refractory multiple myeloma, as established for idecabtagene vicleucel and ciltacabtagene autoleucel. ^1,2^ These responses are not uniformly durable. Relapse can accompany antigen escape, tumour-intrinsic adaptation, limited CAR-T persistence or a suppressive host immune environment. ^3,4^ The early post-infusion period is therefore both a phase of tumour clearance and a period in which lymphodepletion, inflammatory signals, residual malignant plasma cells and recovering host immunity reconstruct the marrow ecosystem.

Single-cell studies have begun to resolve this reconstruction. The source study analysed here identified pre-existing CD39-expressing monocytes and impaired lymphocyte function in nonresponders, alongside heterogeneous CAR-T expansion and exhaustion. ^3^ An independent marrow study linked shorter progression-free survival to BAFF^+^PD-L1^+^ myeloid cells, whereas longer remission was associated with CLEC9A^+^ conventional type 1 dendritic cells and CD27^+^TCF1^+^ T cells. ^5^ A second paired-marrow study reported that favourable responses at day 28 coincided with enhanced cDC1 antigen presentation, stronger CD8 effector activity and reduced regulatory-T-cell immunosuppression.^6^ Together, these observations indicate that treatment outcome reflects coordinated recovery across myeloid and lymphoid compartments rather than CAR-T phenotype alone.

The interpretation of C1Q-expressing macrophages remains less settled. C1Q-associated macrophages recur across human cancers, but tumour-infiltrating myeloid states are heterogeneous and cannot be reduced reliably to an M1–M2 axis.^7,8^ In primary refractory diffuse large B-cell lymphoma, C1QB-expressing macrophages were more abundant in progressive disease, and macrophage cholesterol efflux inhibited CAR-T cytotoxicity in experimental co-culture while inducing a dysfunctional CD8 T-cell state.^9^ In the same study, C1QB macrophages from patients with complete response instead expressed antigen-processing, interferon-γ-response and T-cell-activation programmes. This context dependence raises a narrower possibility: C1Q expression identifies a myeloid remodelling scaffold whose functional meaning depends on the programmes expressed alongside it.

We therefore narrowed the question from a putative cross-cancer C1Q–cholesterol checkpoint to an early, disease-specific hypothesis. After BCMA CAR-T therapy in multiple myeloma, response may accompany an adaptive C1Q-associated myeloid state in which antigen presentation and interferon signalling outweigh cholesterol efflux, TGF-β and hypoxia. We tested this hypothesis by patient-level pseudobulk reanalysis of the publicly available GSE234261 cohort, separating peripheral blood from bone marrow and distinguishing pathway enrichment from patient-level association.

## RESULTS

### Patient-level reanalysis resolves early blood and marrow myeloid states

The public GSE234261 gene-expression matrices comprised 25 specimens from ten patients, including pretreatment peripheral blood and approximately day-30 peripheral-blood or bone-marrow samples. ^3,10^ After sample-wise quality control and doublet filtering, 171,971 of 179,876 input transcriptomes were retained. Broad immune annotation assigned 86,830 cells to the myeloid compartment and 31,653 to CD8 T cells. Counts were aggregated by patient, specimen and compartment before inference; no cell was treated as an independent biological replicate.

The primary comparisons were day-30 complete response (CR) versus nonCR, analysed separately in peripheral blood (PB; nine patients) and bone marrow (BM; six patients). A longitudinal PB comparison included nine paired patients. Six locked programmes were used to test the focused hypothesis: C1Q (*C1QA, C1QB* and *C1QC*), cholesterol efflux, antigen processing and presentation, interferon-γ response, TGF-β response and hypoxia. The three-gene C1Q set was retained as a deliberately narrow marker programme; its enrichment statistics should not be interpreted as evidence for a discrete macrophage lineage.

### Responding blood shows a C1Q-associated antigen-presenting and interferon-rich programme

At day 30 in PB, responding patients showed enrichment of the C1Q programme relative to nonresponders (normalized enrichment score (NES) = 1.70, within-contrast false discovery rate (FDR) = 1.92 × 10^−5^). Antigen processing and presentation was also enriched (NES = 1.99, FDR = 3.96 × 10^−3^), with a particularly strong interferon-γ response (NES = 3.17, FDR = 2.56 × 10^−30^; **Fig. 1**). By contrast, TGF-β signalling was depleted in responders (NES = −1.76, FDR = 3.96 × 10^−3^). Hypoxia trended in the same negative direction but did not meet the within-contrast FDR threshold (NES = −1.22, FDR = 0.119).

**Fig. 1.**
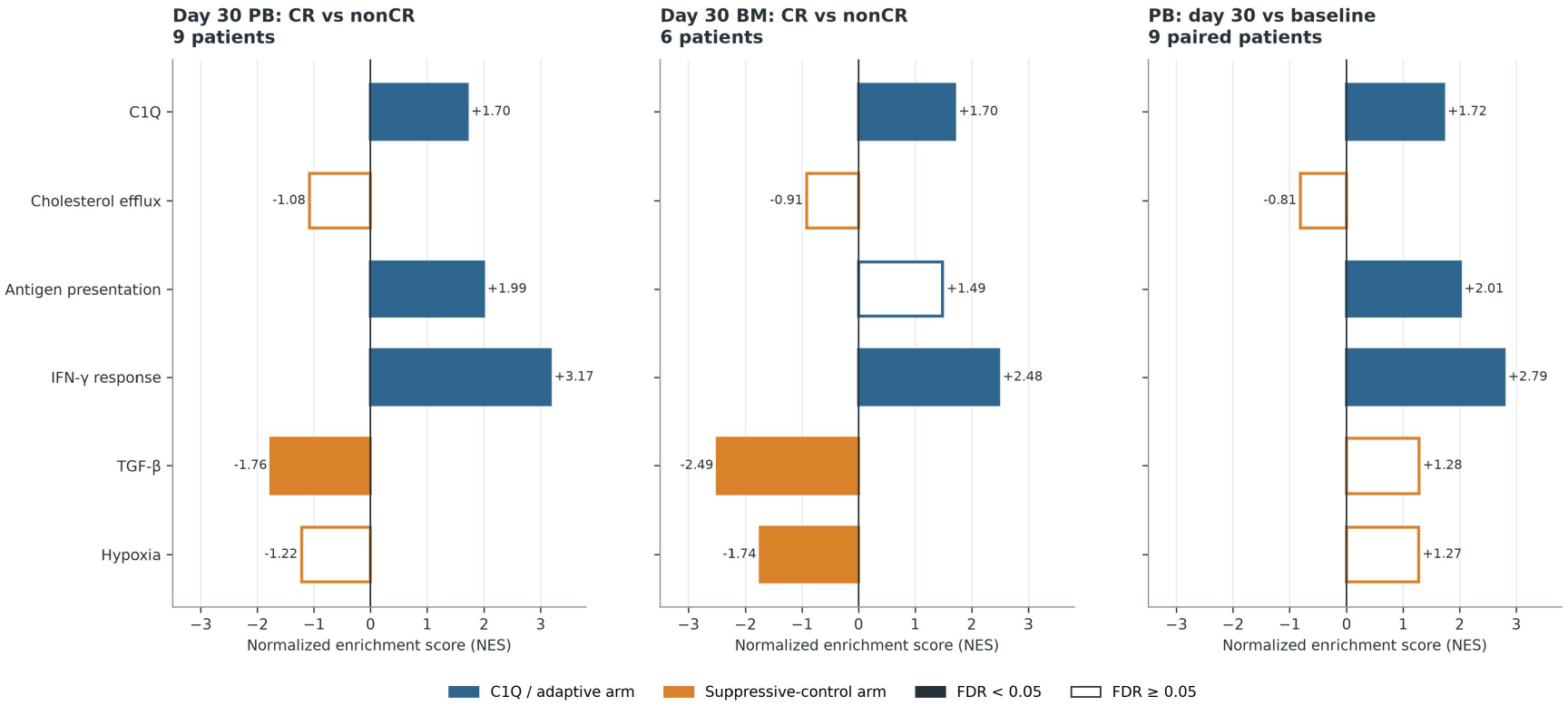
C1Q-associated adaptive myeloid programmes at day 30. Patient-level myeloid pseudobulk gene-set enrichment in GSE234261. The first two panels compare CR with nonCR at day 30 in PB and BM; the third compares day 30 with baseline in paired PB samples. All six locked programmes are shown. Filled bars denote FDR < 0.05 within the displayed contrast, open bars denote FDR ≥ 0.05. Positive NES denotes enrichment in CR or at day 30. PB and BM were analysed independently.

Cholesterol efflux was not enriched in responding PB (NES = −1.08, FDR = 0.365). Thus, the dominant day-30 response-associated pattern was not simultaneous activation of C1Q and cholesterol export. Instead, C1Q co-occurred with antigen presentation and interferon signalling while a canonical suppressive programme was reduced.

### Responding marrow recapitulates C1Q and interferon activation while suppressive programmes decline

Day-30 BM showed a concordant C1Q signal (NES = 1.70, FDR = 1.33 × 10^−4^) and interferon-γ enrichment (NES = 2.48, FDR = 9.08 × 10^−14^; Fig. 1). Antigen presentation was directionally increased but did not pass correction in the six-patient comparison (NES = 1.49, FDR = 0.0838). TGF-β (NES = −2.49, FDR = 1.54 × 10^−7^) and hypoxia (NES = −1.74, FDR = 6.93 × 10^−4^) were depleted in responders. Cholesterol efflux again showed no evidence of positive enrichment (NES = −0.91, FDR = 0.586).

The shared PB–BM features were therefore C1Q and interferon activation, not cholesterol efflux. Tissue-specific differences remained: antigen presentation reached the correction threshold in PB but not BM, whereas depletion of hypoxia was strongest in BM. We consequently did not pool PB and BM or assign a common effect size across tissues.

### Longitudinal blood analysis supports treatment-associated adaptive remodelling

In paired PB samples, C1Q increased from baseline to day 30 (NES = 1.72, FDR = 3.57 × 10^−5^), together with antigen presentation (NES = 2.01, FDR = 9.25 × 10^−4^) and interferon-γ response (NES = 2.79, FDR = 1.72 × 10^−19^; **Fig. 1**). Cholesterol efflux was not enriched (NES = −0.81, FDR = 0.745).

TGF-β and hypoxia were directionally positive in the paired treatment contrast but did not reach the correction threshold (FDR = 0.153 and 0.0804, respectively).

This longitudinal result indicates that the adaptive C1Q-associated programme was not solely a cross-sectional feature of the response groups. However, treatment-associated change and response discrimination are different estimands. The paired analysis tests remodelling after therapy, whereas the CR–nonCR comparison tests whether the resulting state differs with clinical response.

### Patient-level composite scores remain exploratory

Pathway enrichment does not by itself establish a patient-level biomarker. We therefore assessed prespecified sample-level scores. In day-30 BM, the myeloid C1Q–cholesterol composite correlated with CD8 dysfunction (Spearman ρ = 0.943, permutation P = 0.0168; six patients), but the association did not survive correction across the primary patient-level testing family (FDR = 0.247; Fig. 2). The corresponding PB association was weak (ρ = 0.167, P = 0.679, FDR = 0.821; nine patients).

**Fig. 2.**
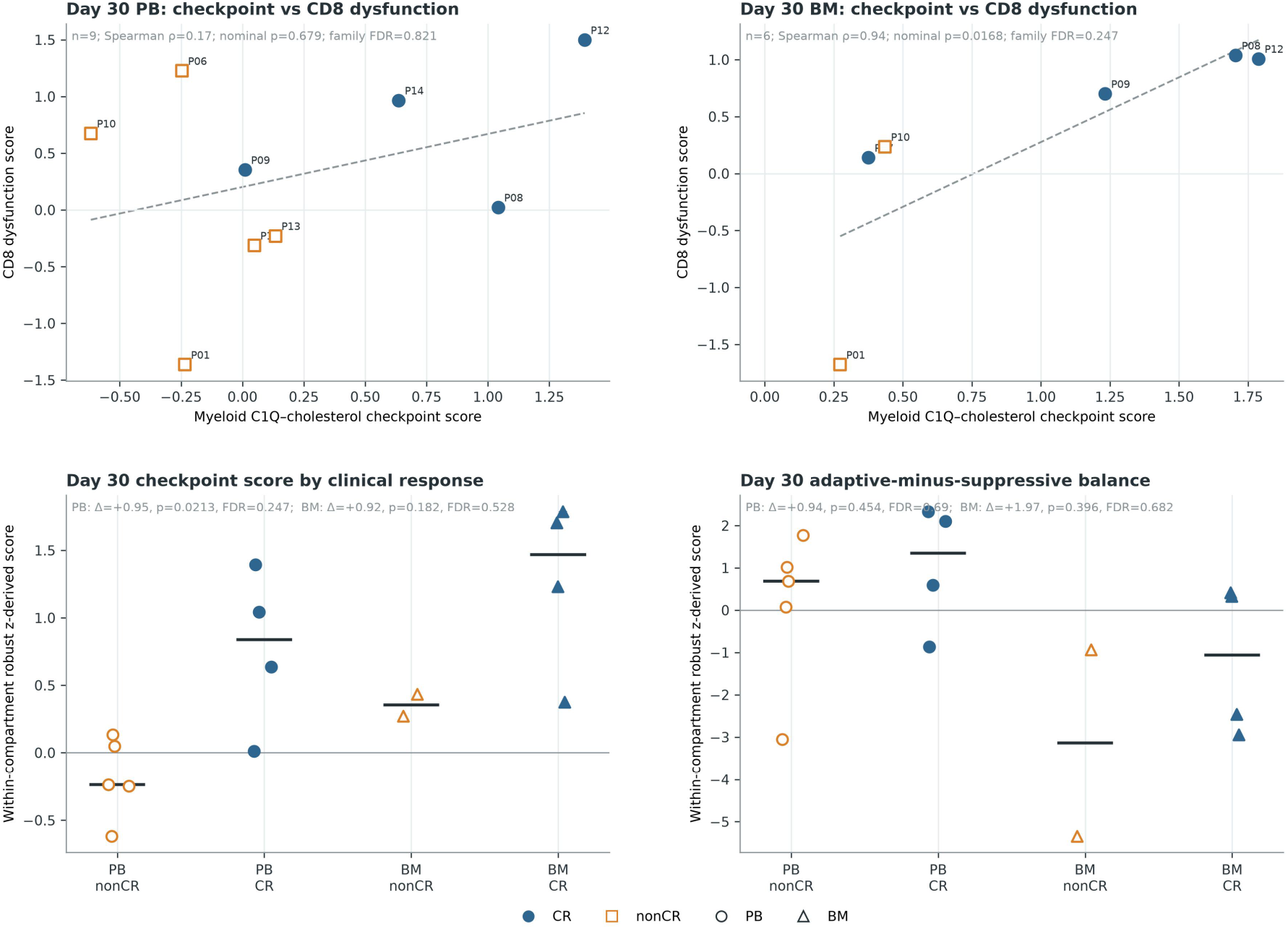
Patient-level associations are compartment-specific and exploratory. Each mark denotes one patient. The upper panels show associations between the myeloid C1Q–cholesterol composite and CD8 dysfunction in PB and BM. Dashed lines are visual linear fits; inference used Spearman rank correlation with 10,000 patient-label permutations. The lower panels show individual checkpoint and adaptive-minus-suppressive scores by response and tissue, with horizontal lines denoting medians. Nominal P values and study-wide patient-level FDR values are displayed to separate effect direction from corrected evidence.

The composite score was higher in CR than nonCR at day 30 in PB (mean standardized difference = 0.954, permutation P = 0.0213, FDR = 0.247) and BM (difference = 0.921, P = 0.182, FDR = 0.528). This direction is opposite to a model in which a larger C1Q–cholesterol score is uniformly suppressive. The adaptive-minus-suppressive balance was also directionally higher in responders in PB (difference = 0.942, P = 0.454, FDR = 0.690) and BM (difference = 1.97, P = 0.396, FDR = 0.682), but uncertainty was substantial. Leave-one-patient-out estimates retained the direction of the response-group mean differences, yet this sensitivity analysis cannot compensate for the small cohort or the absence of an independent outcome-labelled validation dataset.

## DISCUSSION

This reanalysis supports a focused interpretation of early myeloid recovery after BCMA CAR-T therapy. At approximately day 30, response was accompanied by a C1Q-associated programme that aligned with interferon signalling and, most clearly in blood, antigen presentation. In marrow, the same C1Q and interferon signals coincided with depletion of TGF-β and hypoxia programmes. Cholesterol efflux was not positively enriched in responders or after therapy. The data therefore do not support a simple model in which C1Q-high myeloid cells are intrinsically suppressive. They instead favour a state-contingent model: C1Q identifies a macrophage or monocyte remodelling axis whose functional consequence depends on the programmes expressed alongside it.

This interpretation reconciles observations that might otherwise appear contradictory. Pan-cancer atlases show that myeloid phenotypes recur across tissues but remain plastic, and that classical polarization labels obscure functionally distinct macrophage states. ^7,8^ In DLBCL, C1QB macrophages were associated with progression when cholesterol export and suppressive interactions with CD8 T cells dominated. C1QB macrophages from complete responders instead expressed antigen-processing and interferon-related genes. ^9^ Experimental work in malignant pleural effusion further indicates that macrophage-derived C1Q can contribute to lymphocyte dysfunction while participating in macrophage antigen-presentation programmes. ^11^ C1Q expression alone is therefore insufficient to infer immune activation or suppression.

The MM context strengthens the adaptive interpretation. Independent BCMA CAR-T studies associated durable remission with cDC1 recovery, antigen-presentation capacity, T-cell diversity and reduced suppressive myeloid states.^6,12^ A separate longitudinal single-cell atlas similarly implicated endogenous T-cell dysfunction in relapse.^4^ Our analysis adds a narrower observation: within the early post-treatment myeloid compartment, C1Q can accompany the same antigen-presenting and interferon-rich ecology associated with response. This is a refinement rather than a contradiction of the source study, which emphasized immunosuppressive monocytes present before treatment.^3^

Several features prevent a stronger conclusion. First, only ten patients were available, with nine day-30 PB and six day-30 BM profiles. Very small gene sets, particularly the three-gene C1Q programme, can yield extreme enrichment statistics when their members occupy concordant positions in a ranked list; the result shows coordinated expression, not a newly proven cell type. Second, broad myeloid pseudobulk improves control of patient-level replication but averages monocytes, macrophages and dendritic cells. The analysis therefore cannot localize the programme to a purified C1Q-high macrophage subset. Third, response labels were CR versus nonCR rather than a mature duration-of-response endpoint. The manuscript should not equate early complete response with durable remission.

Fourth, FDR values for GSEA were controlled within each prespecified contrast, whereas patient-level permutation tests were corrected across the broader primary testing family. The strong pathway results and the nonsignificant patient-level corrected associations are not inconsistent: they answer different questions, but together they show that transcriptional coherence currently exceeds biomarker certainty. Fifth, the reference MM cohorts lacked a closed, directly comparable public mapping between processed expression files and the same CR–nonCR endpoint; their longitudinal results were therefore not used as independent clinical validation. Finally, the analysis is observational and cannot determine whether the C1Q-associated state promotes response, arises because tumour burden falls, or reflects a shared consequence of treatment exposure.

The next experiment should test the state rather than C1Q alone. A prospective cohort should collect paired baseline and day-28 to day-35 PB and BM, quantify myeloid C1Q protein together with antigen-presentation, interferon, cholesterol-efflux, TGF-β and hypoxia modules, and relate these measurements to minimal residual disease and progression-free survival. Spatial or protein-resolved assays should determine whether the adaptive programme resides in macrophages, cDC1-like cells or mixed myeloid niches. Perturbation of cholesterol export or interferon signalling should be reserved for models in which the relevant state has first been localized and reproduced. Until then, adaptive C1Q-associated remodelling is best treated as a falsifiable ecological hypothesis, not a therapeutic target or validated biomarker.

## METHODS

### Study design and public data

This was a secondary analysis of publicly available human single-cell RNA-sequencing data. GSE234261 contains peripheral-blood mononuclear cells and bone-marrow mononuclear cells obtained before and approximately 30 days after approved BCMA-directed CAR-T therapy. ^3,10^ Processed count matrices were downloaded fresh from GEO. Patient, tissue, time-point and response mappings were derived from GEO metadata and the published study. The analysis was restricted to the 25 deposited gene-expression specimens from ten patients. PB and BM were not pooled.

The focused hypothesis and gene programmes were frozen before the response-directed synthesis. The primary estimand was the day-30 CR–nonCR difference in myeloid transcriptional programmes, separately in PB and BM. The paired baseline-to-day-30 PB contrast was secondary. Correlation between myeloid and CD8 programmes and sample-level response comparisons were exploratory.

### Quality control and immune annotation

Matrices were processed independently by sample. Cells expressing fewer than 200 genes or with more than 20% mitochondrial counts were removed. Doublets were estimated with Scrublet using an expected doublet rate of 0.06. ^13^ Counts from retained cells were preserved for pseudobulk analysis. Normalized log-expression profiles were annotated with the publicly distributed CellTypist `Immune_All_Low` model, followed by deterministic mapping to broad myeloid and CD8 T-cell compartments. ^14^ The inferential unit was the patient or patient–time-point; cell numbers were not used as replicate counts.

### Locked gene programmes

The C1Q programme contained *C1QA, C1QB* and *C1QC*. Cholesterol efflux was defined from GO:0033344. Antigen processing and presentation used a locked public ontology-derived set. Interferon-γ response, TGF-β signalling and hypoxia used public MSigDB Hallmark collections. ^15^ A programme with eight or fewer genes was considered measurable only when every gene was observed; larger programmes required at least three genes and at least 50% coverage. Missing components were not reweighted.

For sample-level summaries, counts per million were log^2^-transformed after adding one, programme genes were averaged, and scores were standardized within cohort and analysis compartment using the median and median absolute deviation. The adaptive arm was the mean of antigen-presentation and interferon-γ z-scores. The suppressive arm was the mean of cholesterol-efflux, TGF-β and hypoxia z-scores. `State balance` was adaptive minus suppressive. The exploratory checkpoint composite was the equal-weight mean of C1Q and cholesterol-efflux z-scores.

### Patient-level pseudobulk differential expression and enrichment

Raw counts were summed by patient, specimen and broad immune compartment. Differential expression used edgeR with trimmed mean of M-values normalization and robust quasi-likelihood generalized linear models. ^16^ Response contrasts modelled CR versus nonCR. Paired longitudinal models included patient as a blocking factor and tested day 30 versus baseline. Genes were filtered with `filterByExpr`. Ranked statistics were defined as the sign of log fold change multiplied by the square root of the quasi-likelihood F statistic.

Gene-set enrichment used a multilevel GSEA implementation with minimum and maximum set sizes of 3 and 500, respectively, following the ranked-list framework. ^17^ FDR was controlled by the Benjamini– Hochberg procedure across the locked programmes within each contrast. Because each contrast used a restricted, prespecified programme family, these FDR values do not control across every contrast in the manuscript.

### Patient-level association tests

Spearman correlations and group mean differences were evaluated with 10,000 patient-label permutations. Paired changes used 10,000 random sign flips. Patient-level P values were adjusted across the primary test family using the Benjamini–Hochberg method. Leave-one-patient-out estimates assessed direction stability; they were not treated as independent validation. All tests were two-sided. No power calculation was performed because the analysis reused all eligible public specimens.

### AI-assisted workflow

AI-assisted tools supported hypothesis formulation, public-dataset discovery, workflow orchestration, protocol critique and language editing, following the evidence-guided BioPathfinder framework and the general-purpose Biomni biomedical research architecture. ^18,19^ Raw expression matrices were processed only by deterministic local scripts. All dataset mappings, statistical calculations, figures, references and scientific conclusions were independently reviewed and approved by the accountable human authors. The AI-assisted tools are not authors and bear no responsibility for the work.

## Data availability

The primary dataset is available from NCBI Gene Expression Omnibus under accession GSE234261. Reference datasets discussed in the manuscript are available under GSE210079 and GSE271915. This study generated no new sequencing data.

## Code availability

The principal code implementing the BioPathfinder workflow is publicly available at https://github.com/TeammateDownloadGenshin/BioPathfinder. Analysis-specific scripts, locked programme definitions, patient-level result tables, figure-source tables and the clean-room manifest are retained in the analysis workspace and will be deposited in a public archival repository before formal submission.

## Acknowledgements

We thank the UCLA/CFAR Virology Core Laboratory for providing healthy-donor peripheral blood mononuclear cells and the UCLA Division of Laboratory Animal Medicine for animal care and support.

## Author contributions

S.W. designed the experiments, analysed the data and wrote the manuscript. S.W. and Q.W. developed the software. Y.-R.L. and S.L. conceived and supervised the study, interpreted the results and revised the manuscript. All authors reviewed and approved the final manuscript.

